# Micromanipulation of amyloplasts with optical tweezers in *Arabidopsis* stems

**DOI:** 10.1101/2020.11.12.379289

**Authors:** Yoshinori Abe, Keisuke Meguriya, Takahisa Matsuzaki, Teruki Sugiyama, Hiroshi Y. Yoshikawa, Miyo Terao Morita, Masatsugu Toyota

## Abstract

Intracellular sedimentation of highly dense, starch-filled amyloplasts toward the gravity vector is likely a key initial step for gravity sensing in plants. However, recent live-cell imaging technology revealed that most amyloplasts continuously exhibit dynamic, saltatory movements in the endodermal cells of *Arabidopsis* stems. These complicated movements led to questions about what type of amyloplast movement triggers gravity sensing. Here we show that a confocal microscope equipped with optical tweezers can be a powerful tool to trap and manipulate amyloplasts noninvasively, while simultaneously observing cellular responses such as vacuolar dynamics in living cells. A near-infrared (λ = 1064 nm) laser that was focused into the endodermal cells at 1 mW of laser power attracted and captured amyloplasts at the laser focus. The optical force exerted on the amyloplasts was theoretically estimated to be up to 1 pN. Interestingly, endosomes and trans-Golgi networks were trapped at 30 mW but not at 1 mW, which is probably due to lower refractive indices of these organelles than that of the amyloplasts. Because amyloplasts are in close proximity to vacuolar membranes in endodermal cells, their physical interaction could be visualized in real time. The vacuolar membranes drastically stretched and deformed in response to the manipulated movements of amyloplasts by optical tweezers. Our new method provides deep insights into the biophysical properties of plant organelles in vivo and opens a new avenue for studying gravity-sensing mechanisms in plants.

## Introduction

Plants change their growth patterns in response to a variety of mechanical stimuli such as touch and gravity (Telewski 2006; Morita 2010; Hagihara and Toyota 2020). Gravitropism is the growth response of a plant to gravity, which generally causes roots to grow downward and shoots to grow upward. Although gravitropism has been extensively studied since the days of Charles Darwin (Darwin and Darwin 1880), the early phase of plant gravitropism, gravity sensing, has not been fully elucidated. The most widely accepted model of gravity sensing is the starch–statolith hypothesis, which predicts that sedimentation of densely starch-filled organelles called amyloplasts triggers gravity sensing in plants (Kiss 2000; Su et al. 2017; Nakamura et al. 2019). This hypothesis has been supported by numerous studies using the model plant *Arabidopsis thaliana* (Morita 2010). Sedimentable amyloplasts are found in root columella and shoot endodermal cells of *Arabidopsis*. Amyloplasts settle downward under the influence of gravity and distribute at the bottom of root columella cells (Leitz et al. 2009). Starch-deficient mutants, e.g., *phosphoglucomutase* (*pgm*), possess non-sedimentable amyloplasts with reduced starch content and exhibit weak gravitropism in both roots and shoots (Kiss et al. 1996; Weise and Kiss 1999). *shoot gravitropism* (*sgr*) *2*, *sgr3*, *sgr4,* and *sgr9* mutants also have non-sedimentable amyloplasts in their endodermal cells and their shoot gravitropic responses are attenuated (Kato et al. 2002; Morita et al. 2002; Yano et al. 2003; Nakamura et al. 2011; Alvarez et al. 2016). These data support the idea that the density of amyloplasts is increased by starch accumulation and that their sedimentation triggers gravity sensing in *Arabidopsis*. However, recent live-cell imaging data revealed that amyloplasts in shoot endodermal cells continuously exhibit dynamic, saltatory movements (Nakamura et al. 2015), which is quite different from the static granules (statoliths) sedimenting at the bottom of cells as predicted by the starch–statolith hypothesis (Kiss 2000). Shoot endodermal cells possess a large central vacuole and the amyloplasts are surrounded by a vacuolar membrane. Amyloplast dynamics are, therefore, strongly dependent not only on gravity but also on the internal cellular environment (e.g., vacuoles and actin filaments) (Nakamura et al. 2011). High-density amyloplasts are pulled down to the bottom of cells by gravity but are diffused by their physical interactions with the vacuolar membrane and/or actin filaments in a random fashion.

Centrifugal acceleration creates a hypergravity condition in which sedimentary movement of high-density amyloplasts is more dominant than random diffusive movements, forcing all amyloplasts to the bottom. This makes centrifugation a useful technique to investigate the relationship between amyloplast dynamics and gravity sensing (Toyota et al. 2014). Indeed, the reduced gravitropic sensitivities in *pgm* and *sgr* mutants were restored by centrifugal hypergravity (Fitzelle and Kiss 2001; Toyota et al. 2014; Mori et al. 2016), which correlated with the resumption of amyloplast sedimentation (Fitzelle and Kiss 2001; Toyota et al. 2013b). These studies suggested that intracellular distribution of amyloplasts toward the gravity vector is critical for gravity perception in plants. Since it is still difficult to perform simultaneous imaging of the amyloplast dynamics and their interactions with vacuoles or actin filaments during centrifugation, neither the critical movements triggering gravity sensing nor the subsequent intracellular events have yet been determined.

The optical tweezer is an innovative technology used to remotely trap and three-dimensionally manipulate a single micrometer-sized object in solution with a focused laser beam (Ashkin 1970; Ashkin et al. 1986). If the refractive index (RI) of the target object is higher than that of its surroundings, the object is attracted to the laser focus by the gradient force (Neuman and Block 2004; Hawes et al. 2010). This optical method has been extensively applied in various biological studies, including plant science (Sparkes 2018). Ashkin and Dziedzic (1989) also utilized optical tweezers to analyze the cytosolic organization of organelles in plant cells. They observed artificial formation of transvacuolar strands by trapping and displacing plant organelles for the first time. Since then, optical tweezers have successfully been used to trap various organelles such as chloroplasts in *Arabidopsis* protoplasts (Andersson et al. 2007), and Golgi bodies and peroxisomes in tobacco (Sparkes et al. 2009; Gao et al. 2016), which revealed physical interactions between chloroplasts and endoplasmic reticulum (ER), Golgi bodies and ER, and peroxisomes and chloroplasts. Optical tweezers were also used to capture and manipulate statoliths in *Chara* rhizoids (Leitz et al. 1995), and amyloplasts in *Arabidopsis* roots (Leitz et al. 2009). These studies moved statoliths and amyloplasts in gravity-sensing cells and observed their intracellular dynamics before and after switching on the trapping laser. Unfortunately, interactions between amyloplasts and other organelles were not examined because the microscope system provided only bright-field images. Moreover, a possibility that the excessive-power laser might capture other invisible organelles in addition to the target amyloplasts (i.e., by non-selective trapping) was not excluded.

In this study, we designed a new optical setup enabling high-resolution imaging of fluorescent signals while non-invasively trapping and manipulating plant organelles with optical tweezers. We optimized the optical parameters and found a minimal laser power to trap amyloplasts but not the endosomes and trans-Golgi network. The theoretical optical force exerted on a single amyloplast was estimated using electromagnetic theories. Furthermore, this new system visualized interactions between amyloplasts and vacuolar membranes in vivo and provided deep insights into the biophysical properties of plant organelles in shoot endodermal cells.

## Materials and methods

### Plant materials and growth conditions

Surface-sterilized seeds of *Arabidopsis thaliana* (Col-0 accession) were sown on agar plates (1× MS salts, 1% (w/v) sucrose, 0.01% (w/v) myo-inositol, 0.05% (w/v) MES, and 0.5% (w/v) gellan gum; pH 5.8), incubated in darkness for 1 day at 4°C, and cultivated at 22°C in a growth chamber under constant light for 2 weeks. Plants grown on plates were transplanted into soil under 16L/8D day/night cycle at 22°C. Longitudinal sections of inflorescence stems were prepared from the plants 5 days after bolting, as previously reported (Nakamura et al. 2011; Toyota et al. 2013b). The stem segments were placed in glass-bottom dishes filled with growth medium (1× MS salts, 1% (w/v) sucrose, 0.05% (w/v) MES, and 0.1% (w/v) agar; pH 5.1), and fixed with a cover glass to prevent dehydration of the samples.

### Optical setup of a confocal laser scanning microscope and optical tweezers

A continuous wave (CW) laser beam from a Nd:YVO_4_ laser (λ = 1064 nm; BL-106SU-FE, Spectra-Physics) was used as a light source for experiments with optical tweezers. After adjusting the laser power with a half-wave plate and a polarizing prism, the laser beam was introduced to an inverted confocal laser scanning microscope (A1R MP+, Nikon) and focused on samples via an oil immersion objective lens (NA = 1.35; Olympus). Amyloplasts and organelles trapped in the laser focus were horizontally manipulated at up to 3 μm s^−1^ speed by moving a motorized stage that was installed on the same microscope. The manipulated movements of amyloplasts and organelles in the horizontal (x-y) direction are independent of gravity that is perpendicular to the x-y direction. The laser power of the optical tweezers through the objective lens was measured using a highly sensitive thermal sensor (12A-V1-ROHS, Ophir) with a laser power meter display (VEGA, Ophir).

Fluorescence and bright-field images of the samples were acquired every 2.1 s using other CW lasers with a wavelength of 488 nm or 561 nm. GFP/Venus and mRFP were excited with 488 and 561 nm lasers, respectively, and the resultant fluorescence signals were obtained with 525/50 nm and 595/50 nm filters, respectively. Autofluorescence (680 nm) of amyloplasts, which was excited with 488 and 561 nm lasers, was acquired with a 700/75 nm filter.

### Calculation of diffusion constants of amyloplasts

To acquire the diffusion constant of a single amyloplast, we analyzed the position of each amyloplast over time by detecting its autofluorescence with an automatic tracking module of an image analysis software (NIS-Elements AR analysis, Nikon). Using these tracing data, the two-dimensional mean square displacement (MSD) of each amyloplast was obtained as previously described (Kusumi et al. 1993; Toyota et al. 2013b). The MSD-Δ*t* plot is linear with a slope of 4D:

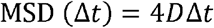

where *D* and Δ*t* are the diffusion constant and time interval, respectively. We acquired the diffusion constant from the slope of the MSD-Δ*t* plot calculated by least-squares methods. With increasing Δ*t*, the number of data used for the calculation of MSD were decreased. To avoid errors caused by using too little data for average calculation, we used a quarter of the total trajectory data in calculations, as previously reported (Saxton and Jacobson 1997; Ruthardt et al. 2011).

### Fluorescence analysis of organellar markers

The organellar markers Ara6-mRFP (Ebine et al. 2011), mRFP-Ara7 (Ueda et al. 2004), Venus-SYP41 (Uemura et al. 2004), and GFP-γTIP (Saito et al. 2005) were expressed under the control of the endodermis-specific SCARECROW promoter (Saito et al. 2005) in this study. The shapes of these organelles are highly dynamic due to their transformative characteristics, making it hard to track single fluorescent spots and calculate the diffusion constants. Thus, we instead used the fluorescence intensity of organelle markers around the laser focus as a parameter to determine whether the organelles were attracted and trapped during laser irradiation. We set a region of interest with a diameter of 3 μm at the center of the laser focus and measured its fluorescence intensity 60 s after the onset of laser irradiation. We calculated the corrected fluorescence intensity (AU) by subtracting the fluorescence intensity at 60 s from the averaged fluorescence intensity at 30 s before laser irradiation.

### Theoretical estimation of optical force exerted on an amyloplast

For calculating the optical force exerted on amyloplasts with size comparable to the wavelength of the trapping laser, complete electromagnetic theories were required to supply an accurate description (Neuman and Block 2004). Hence, we used a computational toolbox for quantitative calculation of optical tweezers based on electromagnetic theory (Nieminen et al. 2007) with the experimental parameters in the software of MATLAB R2019b (MathWorks) as follows: RI of the medium = 1.36; RI of the amyloplast = 1.46; wavelength of laser = 1064 nm; radius of the amyloplast = 1.85 μm; NA of the objective lens = 1.35; and laser power = 0, 0.5, 1, 2, 4, 8, 16, and 30 mW. Note that, since the RI of the amyloplast was not directly measured, we used the median number, 1.46, based on the assumption that it is located between that of chloroplasts (RI = 1.42) and starch granules (RI = 1.48–1.53) (Charney and Brackett 1961; Wolf MJ 1962).

### Statistical analysis

Diffusion constants of single amyloplasts under various laser powers were statistically analyzed by Mann–Whitney *U* tests because these data do not follow a normal distribution. Quantitative measurements of Ara6-mRFP, mRFP-Ara7, and Venus-SYP41 fluorescence were statistically analyzed by Welch’s *t*-test because some pair data exhibited unequal variances. GraphPad Prism (GraphPad software) was used for statistical analysis. All bar graphs are displayed as mean ± SEM.

## Results

### Confocal laser scanning microscope equipped with optical tweezers

The laser was refracted at the boundary between a solvent and a particle with different RIs, where the particle was subjected to the repulsive forces caused by the refraction of the laser and attracted to the laser focus (Figure 1A). We combined the optical tweezers system with a confocal laser scanning microscope to acquire high-resolution bright-field and fluorescent images while remotely manipulating microparticles (Figure 1B). Specimens were irradiated with a near-infrared trapping laser (λ = 1064 nm) (L1, Figure 1B) and excitation lasers (488 nm and 561 nm) (L2, Figure 1B) via an objective lens (Obj, Figure 1B) in the inverted confocal microscope. The resultant bright-field and confocal fluorescent images were acquired with two detectors, D1 and D2, respectively (Figure 1B). To evaluate this novel microscope system, we investigated whether polystyrene beads (RI = 1.57) (Sultanova et al. 2009) in water (RI = 1.33) (Hale and Querry 1973) could be trapped. When the trapping laser was switched on, the beads quickly moved toward the center of the field of view, i.e., the laser focus (Figure 1C; Supplementary Movie S1), indicating that this optical setup is capable of trapping a micro-sized object with a higher RI than the surroundings while simultaneously acquiring images at 70 ms intervals.

**Figure 1.**
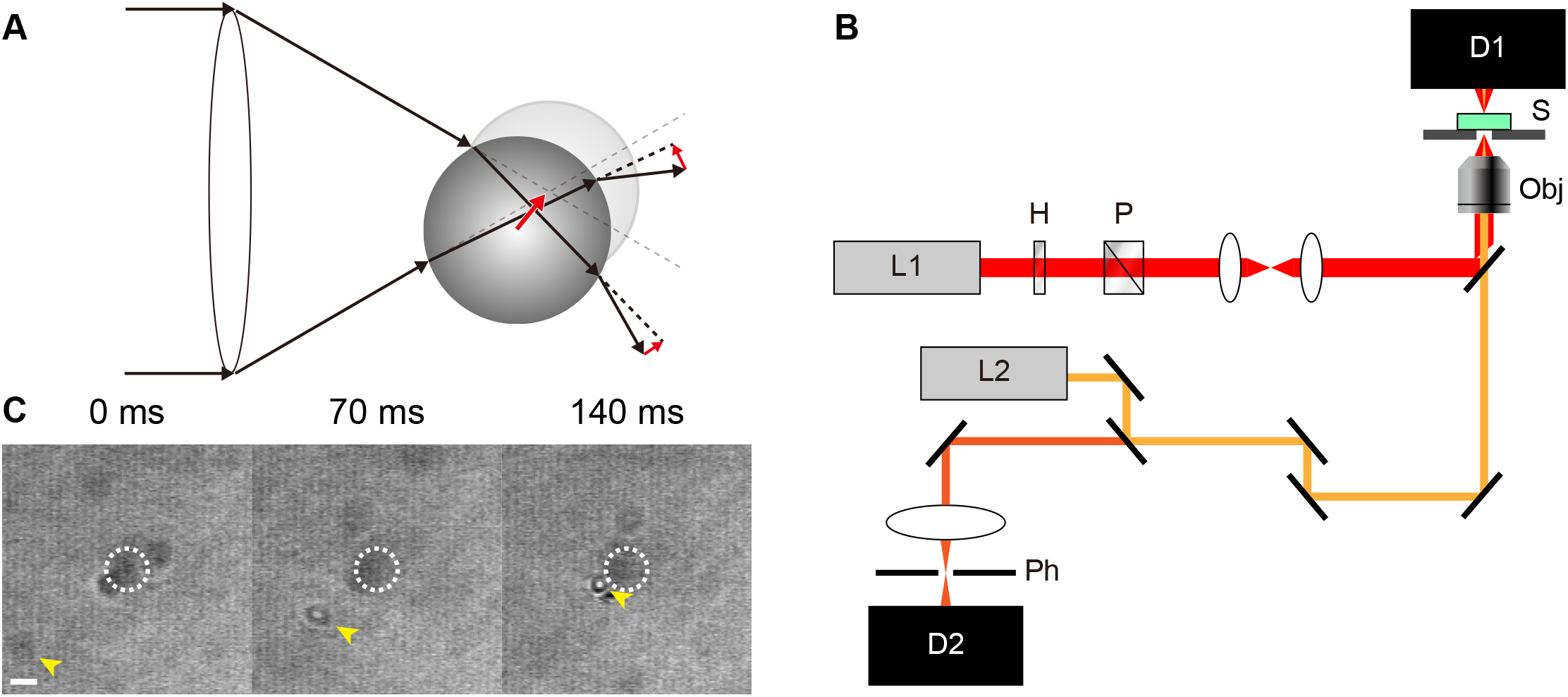
Optical setup of optical tweezers and confocal laser scanning microscope. **A**. Diagram of optical tweezers. Laser beams (black arrows) passing through an objective lens (white ellipse) are refracted at the boundary between the solvent and the particle, which generates optical forces (red arrows) on the particle toward the laser focus (intersection of dashed black lines). **B**. Schematic illustration of the optical setup. Trapping laser (L1) passes through the half-wave plate (H), polarizing prism (P), and objective lens (Obj) to focus on the sample (S). Excitation laser (L2) passes through the same objective lens (Obj) and is focused on the sample (S). Bright-field and fluorescence images are acquired by a detector (D1) and a detector (D2) with a pinhole (Ph), respectively. **C**. Beads’ movements during trapping laser irradiation at 30 mW (yellow arrowheads). Dashed circles represent laser focus of trapping laser. Scale = 2 μm.

### Micromanipulation of the amyloplasts with optical tweezers

The plant organelle amyloplast is filled with starch and plays a crucial role in gravity sensing in the endodermal cells of *Arabidopsis* inflorescence stems (Figure 2A). Because amyloplasts in *Arabidopsis* root columella cells were trapped with optical tweezers (Leitz et al. 2009), their RI is most likely higher than that of the cytoplasm due to their starch content. Thus, we examined whether amyloplasts in the shoot endodermal cells could be trapped and manipulated with the optical tweezers in our new microscope system (Figure 2B). Before turning on the trapping laser, amyloplasts moved randomly from their original position, which resembled the Brownian motion (Figures 2C, D; Supplementary Movie S2). After turning on the trapping laser, amyloplasts were immediately attracted to the laser focus (dashed circles, Figure 2E) and moved along with the position of the laser focus controlled by a motorized stage (Figures 2E, F; Supplementary Movie S2). The trajectories of the amyloplasts before and during irradiation with the trapping laser highlight the different dynamics of the amyloplast movements; the Brownian motion–like dynamics (Figure 2D) was changed into the directional movement by optical tweezers (Figure 2F). These results indicated that our optical system can remotely manipulate amyloplasts while detecting bright-field and confocal fluorescent images.

**Figure 2.**
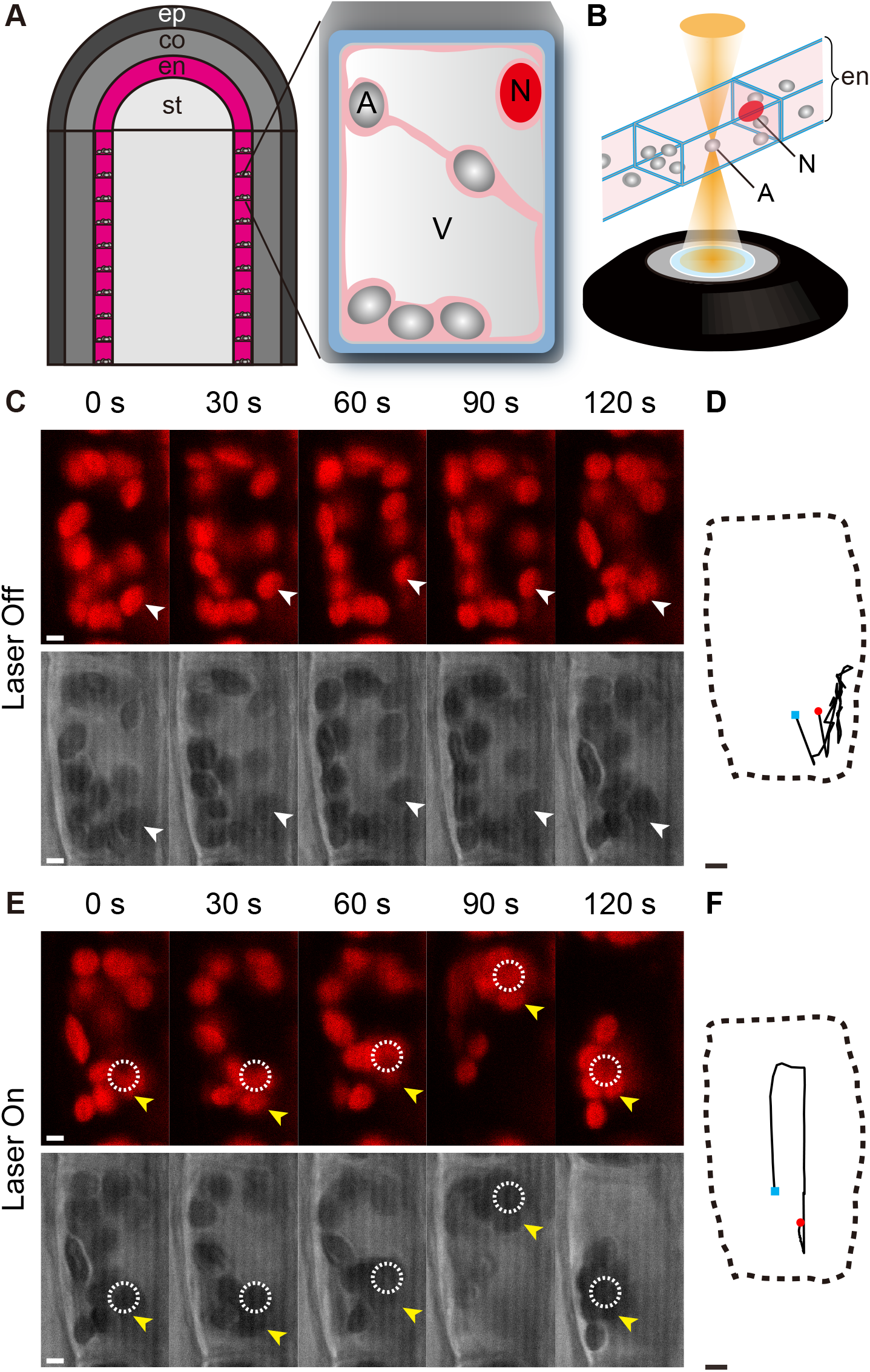
Optical trapping of the amyloplasts in endodermal cells of *Arabidopsis* inflorescence stems. **A**. Diagram of the endodermal cells. ep = epidermis; co = cortex; en = endodermis; st = stele; A = amyloplasts; N = nucleus; V = vacuole. **B**. Schematic illustration of optical trapping of the amyloplasts. **C–F**. Time-lapse images of amyloplast movements and trajectories of the representative amyloplasts before (white arrowheads, C and D) and during laser irradiation (yellow arrowheads, E and F). Autofluorescence of the amyloplasts (upper panels) and bright-field images (lower panels) are shown (C and E). Dashed circles denote the laser focus of the trapping laser (E). Dashed lines = outline of the endodermal cell; red and blue dots = the position of the amyloplast at 0 and 120 s, respectively (D and F). Scale = 2 μm.

### Quantitative analysis of the amyloplast dynamics

To gain further insight into amyloplast dynamics, the relationship between the laser power and the state of amyloplasts (i.e., diffusion constants) was investigated using trajectory data of amyloplasts. When the laser power was set to the minimum non-detectable level (~0 mW), amyloplasts continued to move randomly and left the laser focus (Figure 3A, B; Supplementary Movie S3). The diffusion constant of the amyloplasts was 4.4 × 10^−2^ μm^2^ s^−1^ (*n* = 44) (Figure 3G). Interestingly, almost all amyloplast movements were confined around the laser focus at 1 mW (Figures 3C, D; Supplementary Movie S4), and all the amyloplasts were stably trapped at 30 mW (Figure 3E, F; Supplementary Movie S5). The diffusion constants were 1.9 × 10^−2^ μm^2^ s^−1^ (*n* = 8) and 2.3 × 10^−4^ μm^2^ s^−1^ (*n* = 5) at 1 and 30 mW, respectively, which were statistically lower than those at <0.5 mW (Figure 3G). As the microparticles were trapped and their movement suppressed by the optical tweezers, their diffusion constants became smaller. Therefore, we concluded that 1 mW is a minimum and sufficient laser power to trap shoot amyloplasts using our optical setup (Figure 3G).

**Figure 3.**
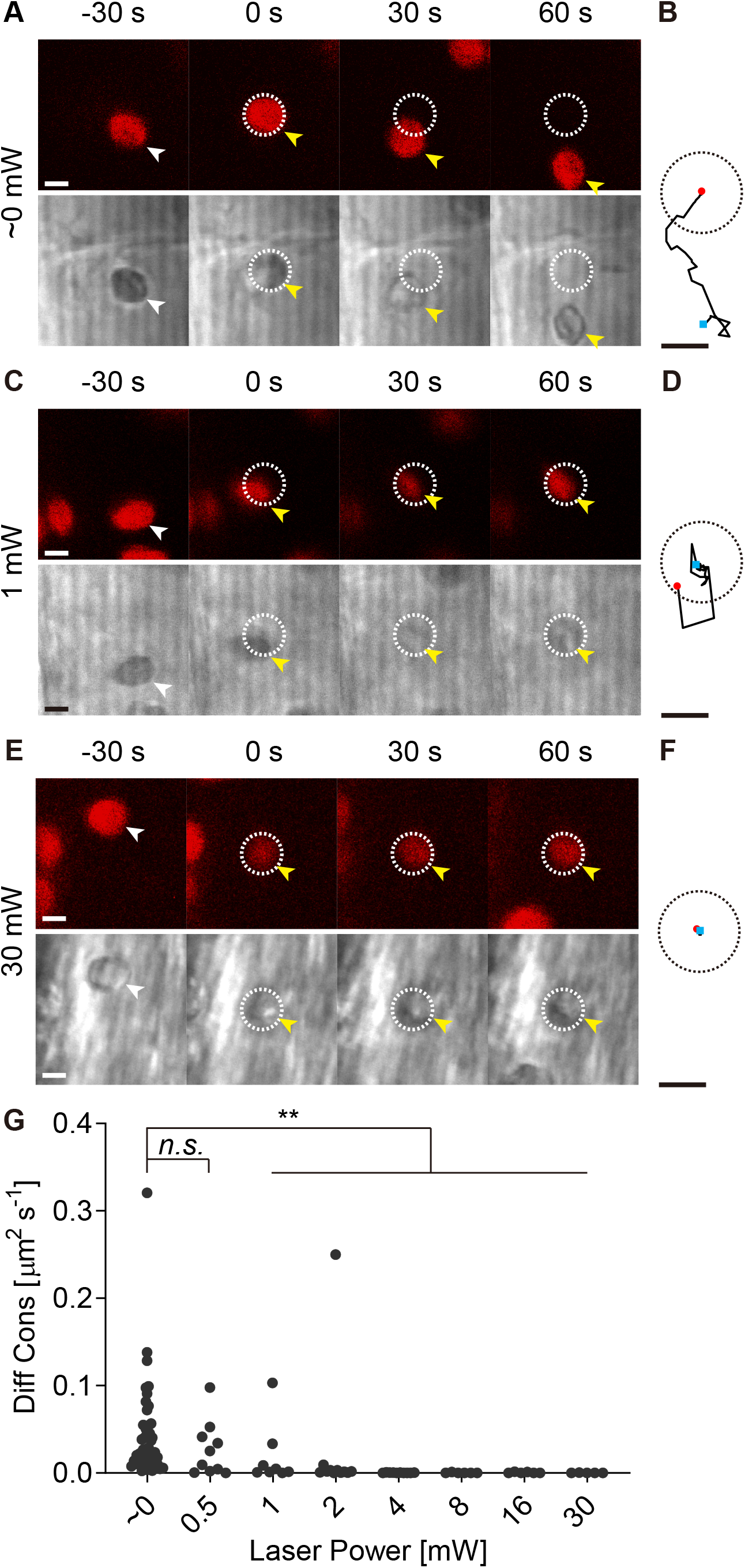
Amyloplast dynamics during trapping laser irradiation. **A–F**. Time-lapse images and trajectories of the representative amyloplasts before (white arrowheads) and during laser irradiation (yellow arrowheads) at ~0 mW (A and B), 1 mW (C and D), and 30 mW (E and F). Autofluorescence of the amyloplasts (upper panels) and bright-field images (lower panels) are shown (A, C, and E). Dashed circle = the laser focus of the trapping laser; red and blue dots = the position of the amyloplast at 0 and 120 s, respectively. Scale = 2 μm. **G**. Distribution of the diffusion constants of amyloplasts when irradiated at various laser powers. *n.s* = not significant*, p* > 0.05; asterisks (*) = significant, *p* < 0.01 (*n* ≥ 5).

### Optical trapping of endosomes and trans-Golgi network

We investigated the dynamics of other organelles during laser irradiation, since the excessive-power laser might trap other organelles in addition to the amyloplasts (known as non-selective trapping). Endosomes and the trans-Golgi network (TGN) are plant organelles critical for intracellular membrane trafficking in plant gravity sensing. Ara6 and Ara7, which belong to Rab/Ypt GTPase family, regulate effector proteins involved in endosomal transport pathways and are localized in two different endosomes, although their expression patterns partially overlap (Ueda et al. 2004; Ueda et al. 2001). SYP41, which is one of SNARE proteins that comprise the Q-SNARE complex in the TGN, regulates the vesicle transport pathway (Uemura et al. 2012). We visualized endosomes and the TGN using their markers (Ara6-mRFP, mRFP-Ara7 and Venus-SYP41, respectively) during laser irradiation at ~0, 1, and 30 mW. No changes in the fluorescence signals of the organelles were observed around the slow-moving laser focus when the trapping laser was powered at ~0 and 1 mW (Figures 4A, 4B, 5A, 5B; Supplementary Figures 1A, 1B; Supplementary Movies S6, S7, S9, S10, S12, S13). In contrast, the organellar fluorescence signals accumulated during laser irradiation at 30 mW and were manipulated by the movement of this laser (Figures 4C, 5C; Supplementary Figure 1C; Supplementary Movies S8, S11, S14). Quantitative analysis of these organellar markers showed that the averaged fluorescence intensities around the laser focus at 30 mW were significantly higher than those at ~0 and 1 mW at 60 s (Figures 4D, 5D; Supplementary Figure 1D). These data indicate that 30 mW, but not 1 mW, is needed to trap the endosomes and TGN in endodermal cells, probably due to lower RIs of these organelles than that of amyloplasts. Taken together with the results of amyloplast trapping (Figure 3), 1 mW lasers can trap only the amyloplasts without affecting the dynamics of other organelles, such as endosomes and the TGN.

**Figure 4.**
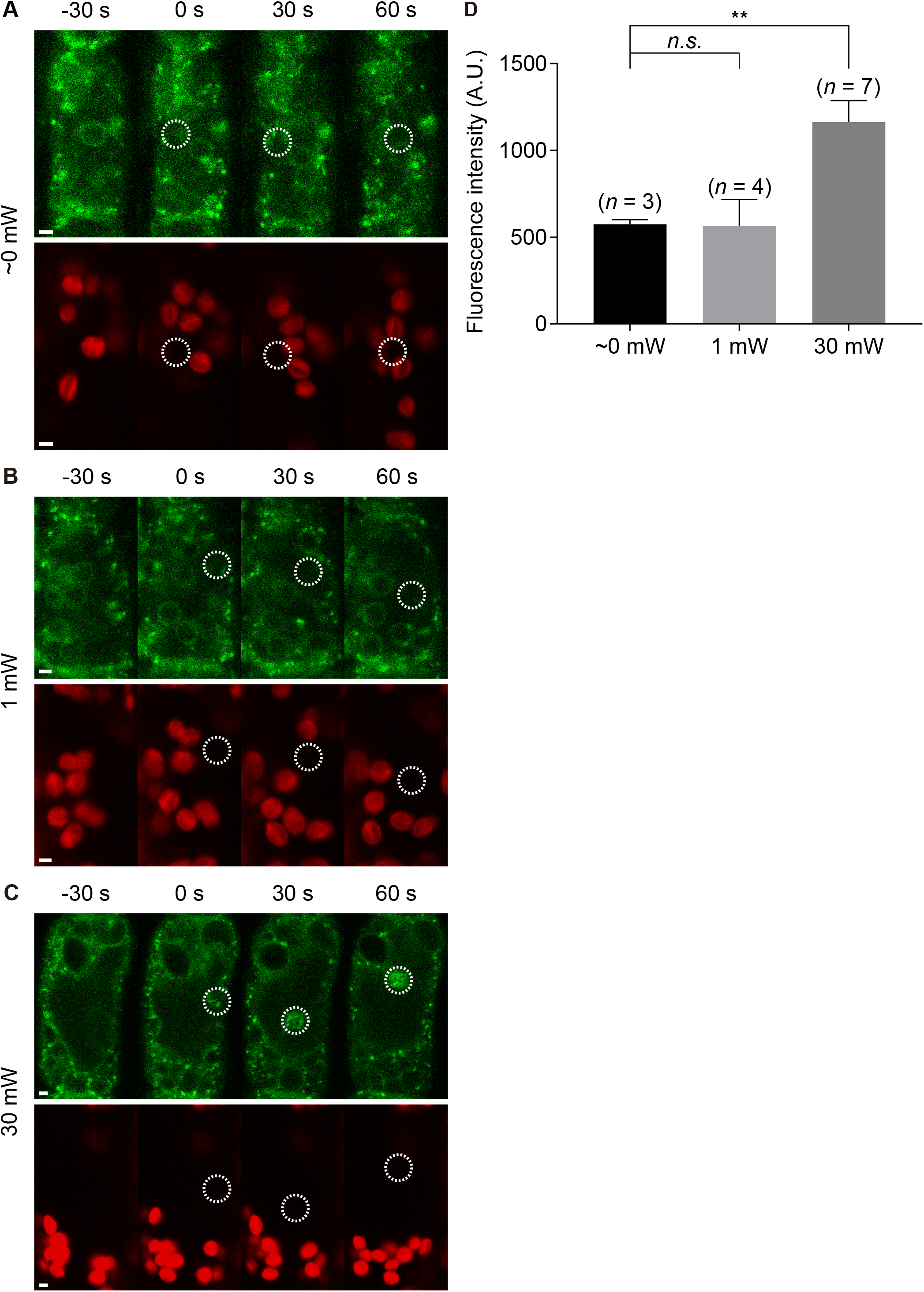
Dynamics of Ara6-mRFP during trapping laser irradiation. **A–C**. Fluorescent images of Ara6-mRFP (upper panels) and autofluorescence images of amyloplasts (lower panels) when irradiated at ~0 mW (A), 1 mW (B), and 30 mW (C). Dashed circles denote the laser focus of the trapping laser. Scale = 2 μm. **D**. Comparison of fluorescence intensities in the dashed circles at 60 s at ~0, 1, and 30 mW. *n.s* = not significant*, p* > 0.05); asterisks (*) = significant, *p* < 0.01.

**Figure 5.**
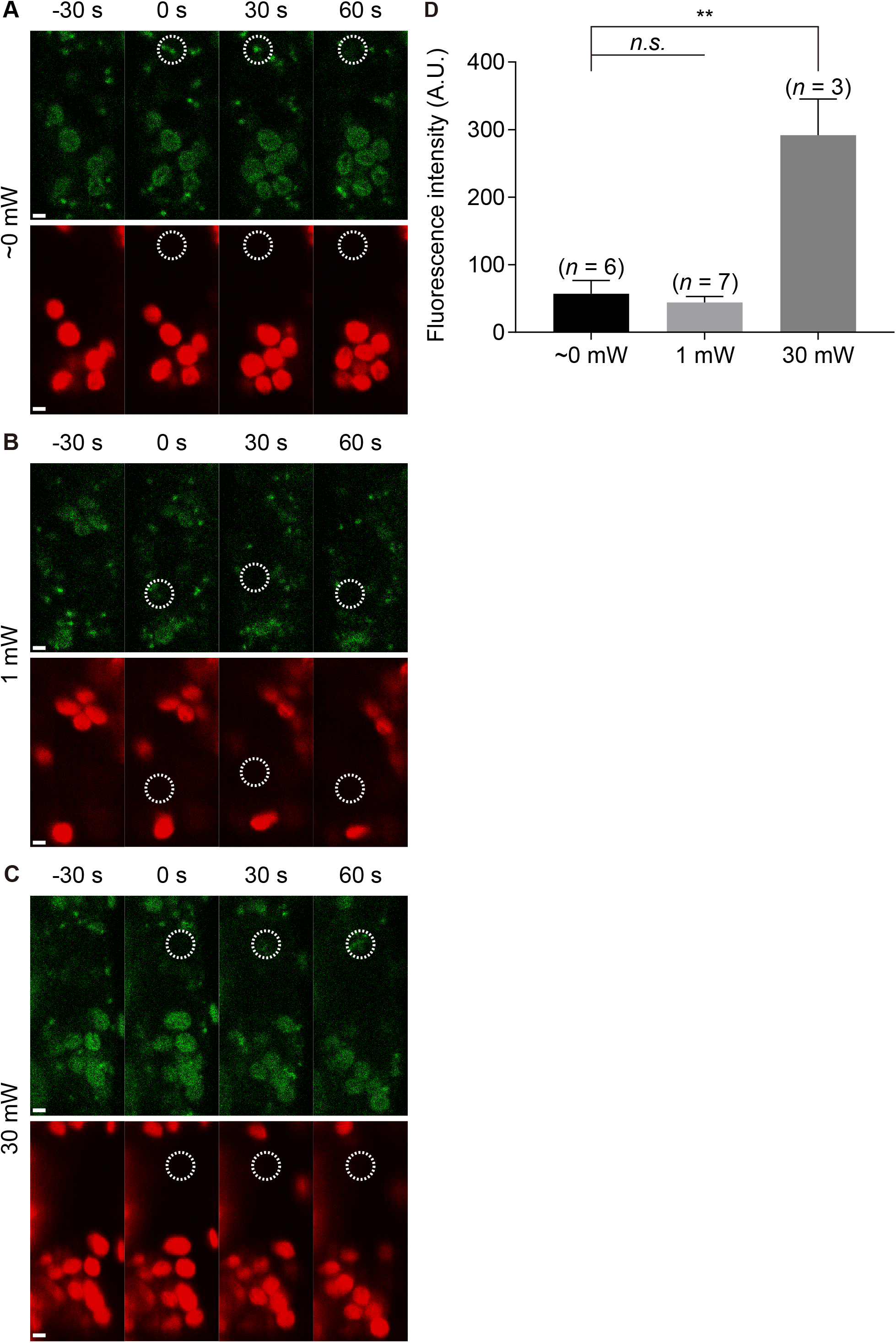
Dynamics of Venus-SYP41 during trapping laser irradiation. **A–C**. Fluorescent images of Venus-SYP41 (upper panels) and autofluorescence images of amyloplasts (lower panels) when irradiated at ~0 mW (A), 1 mW (B), and 30 mW (C). Dashed circles denote the laser focus of the trapping laser. Scale = 2 μm. **D**. Comparison of fluorescence intensities in the dashed circles at 60 s at ~0, 1, and 30 mW. *n.s* = not significant *p* > 0.05; asterisks (*) = significant, *p* < 0.05.

### Optical force exerted on an amyloplast

We estimated the optical force exerted on a single amyloplast at 1 mW in the x-y direction (F_x-y_, Figure 6A) and the optical axis (z) direction (F_z_, Figure 6B) using an optical tweezers computational toolbox (Nieminen et al. 2007). The maximum F_x-y_ and F_z_ were simulated to be approximately 1.0 and 0.7 pN, respectively, when an amyloplast was located around 1.9 μm from the center of the laser focus in the x-y direction (Figure 6A) and 1.7 μm in the optical axis direction (Figure 6B), respectively. Therefore, up to 1 pN is exerted on a single amyloplast, depending on its relative position to the trapping laser. This estimation also illustrated a linear relationship between the laser power and the optical force exerted on an amyloplast (Figure 6C): when an amyloplast was irradiated with the trapping laser at 30 mW (Figure 2), a maximum of 30 pN optical forces worked on it.

**Figure 6.**
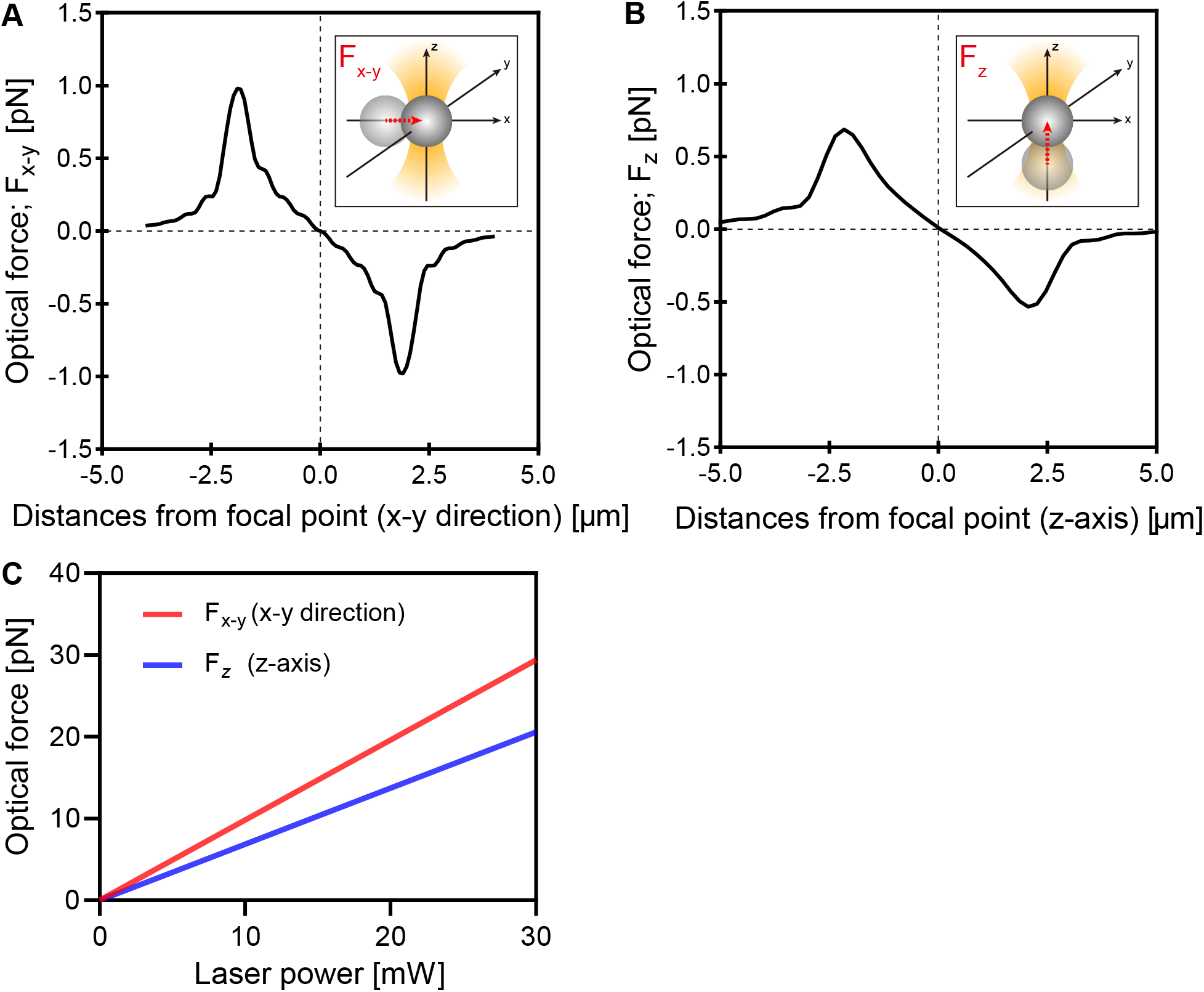
Theoretical estimation of the optical force exerting on a single amyloplast. **A–B**. Optical force in x-y direction (F_x-y_) and optical axis direction (F_z_) at 1 mW laser power. **C**. Relationship between the maximum optical force and the trapping laser power. Red line = F_x-y_; blue line = F_z_.

### Dynamics of vacuolar membranes while manipulating the amyloplasts

In endodermal cells, a central vacuole occupies a large portion of the cell volume and affects amyloplast movement (Figure 2A). To investigate the interactions between vacuolar membranes and amyloplasts, we visualized the vacuole dynamics upon manipulation of the trapped amyloplasts. When we moved the amyloplasts toward the periphery of the cell, the vacuolar membranes drastically stretched and deformed in response (Figure 7A; Supplementary Movie S15). When the trapped amyloplasts were displaced toward the center of the cell, a transvacuolar strand was artificially created (Figure 7B; Supplementary Movie S16). Interestingly, these amyloplasts slowly moved back to their original positions after the trapping laser was switched off (Figure 7C; Supplementary Movie S16). These results suggest that vacuolar membranes possess both elastic and plastic characteristics.

**Figure 7.**
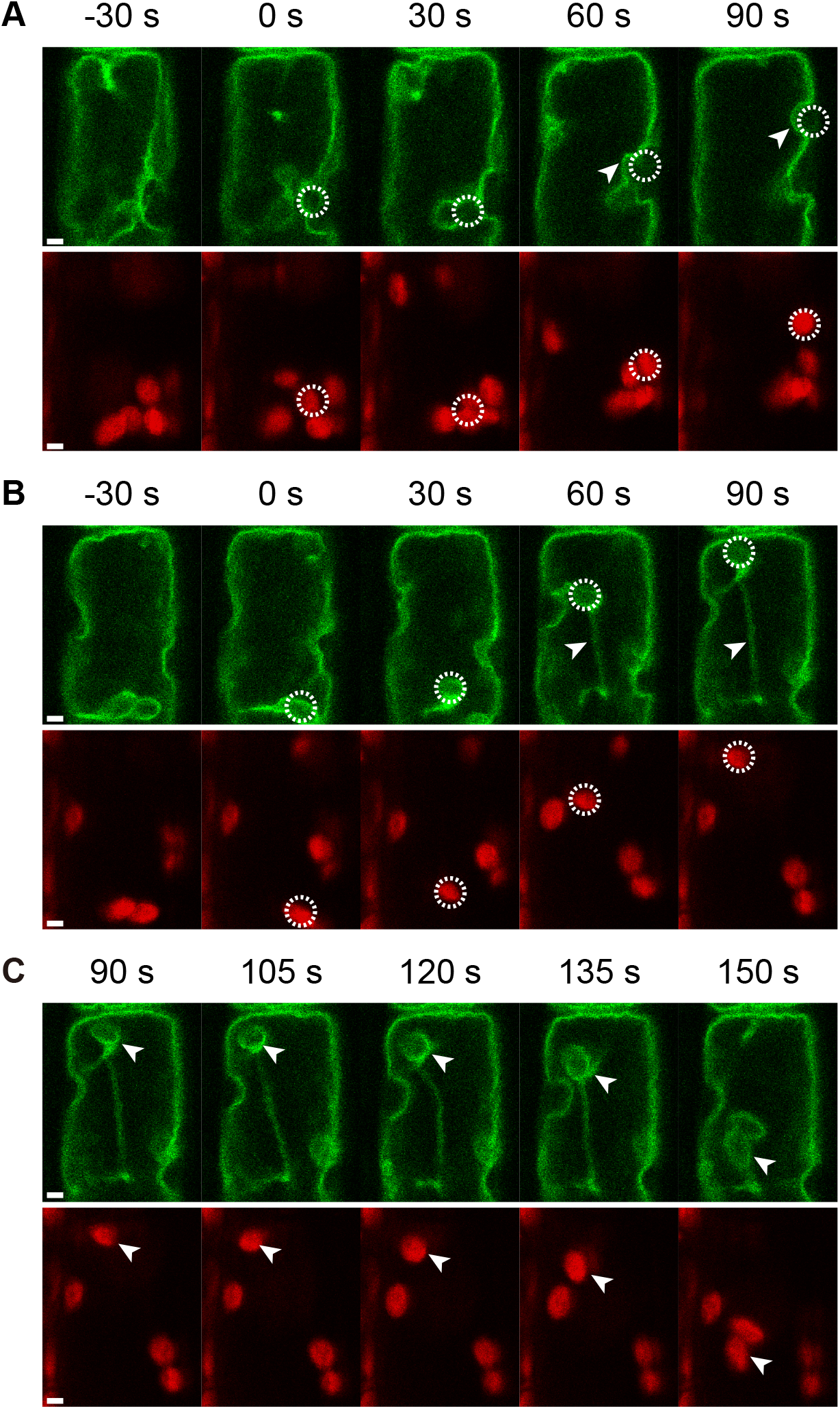
Dynamics of vacuolar membranes and amyloplasts during trapping laser irradiation. **A–C**. Fluorescent images of GFP-γTIP (upper panels) and autofluorescence images of amyloplasts (lower panels) when irradiated at 30 mW. Arrowheads indicate drastic deformation of the vacuolar membranes (A), artificial formation of the transvacuolar strand (B), and “spring back” response of the amyloplasts after switching off the trapping laser (C). Dashed circles denote the laser focus of the trapping laser. Scale = 2 μm.

## Discussion

### Micromanipulation of amyloplasts by optical tweezers: A new method to study gravity sensing in plants

We found that, at 1 mW, our optical tweezers system manipulated the amyloplasts, but not endosomes and the TGN, in endodermal cells (Figure 3), suggesting that this setup may be a powerful tool for research focusing on amyloplast movements and downstream events, namely gravity sensory transduction. Although we did not test the other organelles such as the nucleus, ER, peroxisomes, and mitochondria, it is unlikely that irradiation with the trapping laser at 1 mW would capture them together with amyloplasts. The RI of nuclei in Hela cells is estimated to be 1.35–1.37 (Liu et al. 2016), which is almost identical to that of the cytosol (RI = 1.36). Furthermore, a 300–400 mW high-power laser is required to trap the nucleolus and suppress its intracellular movement in root hair cells of *Arabidopsis* (Ketelaar et al. 2002). The RI of ER in Hela cells is estimated to be 1.37–1.40 (Sandoz et al. 2019), which is slightly higher than that of the cytosol. However, a previous paper reported that ER markers could not be trapped at 15–150 mW in leaf epidermal cells of *Arabidopsis* (Sparkes et al. 2009). While the RI of peroxisome has not yet been reported, 13 mW laser power is needed to trap it in tobacco (Gao et al. 2016). The RI of purified mitochondria is 1.41 (Haseda et al. 2015; Gao et al. 2016), which is higher than that of the cytosol but still lower than that of amyloplasts (RI = 1.46). Furthermore, the size of mitochondria in *Arabidopsis* is 1–3 μm (Logan and Leaver 2000; Haseda et al. 2015), which is smaller than amyloplasts (3.7 μm). Indeed, 30 mW lasers failed to trap mitochondria in the amoeba *Reticulomyxa* (Ashkin et al. 1990). It is most likely that laser irradiation at 1 mW can selectively trap amyloplasts, but not other organelles, in shoot endodermal cells.

### Comparison of optical, gravitational, and endogenous forces

The optical force exerted on a single amyloplast was simulated to be a maximum of 1.0 pN at 1 mW (Figure 6). Gravitational force exerted on an amyloplast (F_AM_) was estimated with the following formula:

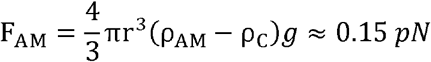

where r is the radius of the amyloplast (1.85 μm), ρ_AM_ is the density of the amyloplast (1.615 × 10^3^ kg m^−3^) (Hinchman and Gordon 1974), ρ_C_ is the density of cytoplasm (1.05 × 10^3^ kg m^−3^) (Hinchman and Gordon 1974), and *g* is the gravitational acceleration (9.8 m s^−2^). The optical force at 1 mW is up to seven times higher than the gravitational force.

We very rarely observed amyloplasts moving away from the laser focus, even at 2 mW (Figure 3G; Supplementary Figure S2; Supplementary Movies S17, S18) where the optical force was up to 2.0 pN (Figure 6C). This phenomenon suggests that intracellular dynamics such as cytoplasmic streaming or transvacuolar strands transiently create more than 2 pN internal force to make amyloplasts escape from the optical force. In this case, the endogenous force is approximately 13 times higher than the gravitational force, which is consistent with recent live-cell imaging data showing that amyloplasts continuously exhibit dynamic, saltatory movements (Nakamura et al. 2015). Furthermore, Berut et al. (2018) reported that amyloplasts were strongly agitated by “intracellular factors” in wheat coleoptiles, but plants can sense the small inclination by using the redistribution of amyloplasts caused by this strong agitation. These results suggest that the intracellular sedimentary movements and dynamic agitations of amyloplasts play an important role in gravity sensing in plants.

Cytoplasmic streaming might be the “intracellular factor” that creates the endogenous force. Given that cytoplasmic streaming creates drag force on an amyloplast (D_AM_) in fluidic liquid, D_AM_ can be calculated with the following formula:

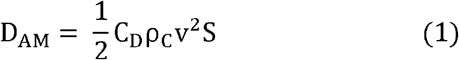

where C_D_ is the drag coefficient, v is the maximum velocity of cytoplasmic streaming (4.0 × 10^−6^ m s^−1^) in leaf epidermal cells of *Arabidopsis* (Tominaga et al. 2013), and S is the area of an amyloplast ((1.85 × 10^−6^)^2^ × π m^2^). The drag coefficient is determined by the Reynolds number, which is a dimensionless number used to predict the flow condition. The Reynolds number for an amyloplast moving in the cytosol is given by:

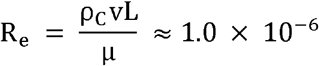

where L is the diameter of an amyloplast (3.7 × 10^−6^ m) and μ is the viscosity of cytosol (0.02 kg m^−1^ s^−1^) (Audus 1962). In this case, R_e_ is 1.0×10^−6^, which is much smaller than 0.5. Therefore, C_D_ can be approximated as follows (Goossens 2019):

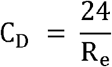

which permits us to rewrite (1) as

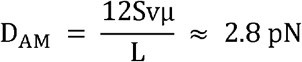

The drag force exerted on an amyloplast is up to 2.8 pN, which is consistent with our observation that amyloplasts very rarely moved away from the 2 pN optical force at 2 mW (Supplementary Figures 2C, D), but not 4 pN at 4 mW (Figure 3G). These estimations support the idea that drag force caused by cytoplasmic streaming plays a critical role in the intracellular factors that diffuse amyloplasts. Cytoplasmic streaming is supposed to be regulated by the sliding motion of the motor protein myosin along actin filaments. Previous papers reported that the dynamic saltatory movements of amyloplasts are attenuated by latrunculin B, an inhibitor of actin polymerization (Nakamura et al. 2011). Actin filaments may mediate intracellular factors via cytoplasmic streaming and/or direct interaction with amyloplasts.

### Interaction between amyloplasts and vacuoles

In endodermal cells, the amyloplasts, which exist in the extremely narrow cytosolic space, are in close proximity to the vacuolar membranes. Therefore, the amyloplast movements appear to be affected by vacuoles. Indeed, abnormal behavior of the vacuolar membrane in *sgr2* and *sgr4* pushes amyloplasts to the periphery of the cell, which restricts amyloplast movement (Kato et al. 2002; Saito et al. 2005). In this study, the vacuole membranes were severely deformed by the manipulated movements of amyloplasts with optical tweezers, resulting in artificial formation of transvacuolar strands (Figures 7A, B). These results suggest that vacuolar membranes possess a high fluidity and plastic characteristics. Interestingly, when the trapping laser was switched off, the amyloplasts returned to their original positions and the transvacuolar strands disappeared (Figure 7C), which resembled a phenomenon called “spring back” (Sparkes 2018). This spring back response was also observed in amyloplasts of root columella cells (Leitz et al. 2009) and tobacco peroxisomes (Gao et al. 2016). The spring back of amyloplasts in shoot endodermal cells may be caused by the elastic properties of vacuolar membranes. However, there is a time lag (approximately 30 s) for the amyloplasts to start moving back to their original position after the laser is switched off (Figure 7C; Supplementary Movie S16). This elastic spring back response may be regulated by more complex interactions with other organelles. Vacuolar membranes affect the amyloplast dynamics and vice versa. This physical interaction and resultant membrane stretch might be involved in gravity responses via mechano-sensitive ion channels (Toyota et al. 2013a; Tatsumi et al. 2014a; Tatsumi et al. 2014b; Iida et al. 2014). Our new system provides novel insights into biophysical characteristics of plant organelles in vivo and a chance to investigate gravity sensory transduction mechanisms in plants.

## Acknowledgements

We thank Yasuko Hashiguchi and Center for Emergent Functional Matter Science of National Chiao Tung University from The Featured Areas Research Center Program within the framework of the Higher Education Sprout Project by the Ministry of Education (MOE) in Taiwan for the technical assistances. This work was supported by Japan Society for the Promotion of Science, KAKENHI (17H05007 and 18H05491 to M.T., 19KK0128 and 20K21117 to H.Y.Y. and JP16H06507 to T.S.); Japan Science and Technology Agency, PRESTO (to M.T.M.) and the Ministry of Science and Technology in Taiwan (MOST 109-2113-M-009-008-to T.S.).

## Supplementary Figure Legends

Supplementary Figure 1. Dynamics of mRFP-Ara7 during trapping laser irradiation. **A–C**. Fluorescent images of mRFP-Ara7 (upper panels) and autofluorescence images of amyloplasts (lower panels) when irradiated at ~0 mW (A), 1 mW (B), and 30 mW (C). Dashed circles denote the laser focus of the trapping laser. Scale = 2 μm. **D**. Comparison of fluorescence intensities in the dashed circles at 60 s at ~0, 1, and 30 mW. *n.s* = not significant *p* > 0.05; asterisks (*) = significant, *p* < 0.001.

Supplementary Figure 2. Amyloplasts that moved away from the laser focus during trapping laser irradiation at 1 and 2 mW. **A–D**. Time-lapse images and trajectories of the representative amyloplasts before (white arrowheads) and during laser irradiation (yellow arrowheads) at 1 mW (A and B) and 2 mW (C and D). Autofluorescence of the amyloplasts (upper panels) and bright-field images (lower panels) are shown (A and C). Dashed circles = the laser focus of the trapping laser; red and blue dots = the position of the amyloplast at 0 and 120 s, respectively. Scale = 2 μm.

## Supplementary Movie Legends

Supplementary Movie 1. Beads’ dynamic before and during trapping laser irradiation (70 ms intervals, total 5 s). Dashed circle and yellow arrow denote the laser focus of the trapping laser and the representative bead moving to the laser focus, respectively.

Supplementary Movie 2. Amyloplasts dynamic before and during trapping laser irradiation at 30 mW (2.1 s intervals, total 344 s). Dashed circles denote the laser focus of the trapping laser. White and yellow arrows indicate amyloplasts before and during laser irradiation, respectively.

Supplementary Movie 3. Amyloplast dynamics during trapping laser irradiation at ~0 mW (2.1 s intervals, total 60 s). Dashed circle denotes the laser focus of the trapping laser. Yellow arrow indicates amyloplasts during laser irradiation.

Supplementary Movie 4. Amyloplast dynamics during trapping laser irradiation at 1 mW (2.1 s intervals, total 60 s). Dashed circles denote the laser focus of the trapping laser. Yellow arrow indicates trapped amyloplasts during laser irradiation.

Supplementary Movie 5. Amyloplast dynamics during trapping laser irradiation at 30 mW (2.1 s intervals, total 60 s). Dashed circles denote the laser focus of the trapping laser. Yellow arrow indicates trapped amyloplasts during laser irradiation.

Supplementary Movie 6. Ara6-mRFP dynamics during trapping laser irradiation at ~0 mW (4.2 s intervals, total 60 s). Dashed circles denote the laser focus of the trapping laser.

Supplementary Movie 7. Ara6-mRFP dynamics during trapping laser irradiation at 1 mW (4.2 s intervals, total 60 s). Dashed circles denote the laser focus of the trapping laser.

Supplementary Movie 8. Ara6-mRFP dynamics during trapping laser irradiation at 30 mW (4.2 s intervals, total 60 s). Dashed circles denote the laser focus of the trapping laser.

Supplementary Movie 9. mRFP-Ara7 dynamics during trapping laser irradiation at ~0 mW (4.2 s intervals, total 60 s). Dashed circles denote the laser focus of the trapping laser.

Supplementary Movie 10. mRFP-Ara7 dynamics during trapping laser irradiation at 1 mW (4.2 s intervals, total 60 s). Dashed circles denote the laser focus of the trapping laser.

Supplementary Movie 11. mRFP-Ara7 dynamics during trapping laser irradiation at 30 mW (4.2 s intervals, total 60 s). Dashed circles denote the laser focus of the trapping laser.

Supplementary Movie 12. Venus-SYP41 dynamics during trapping laser irradiation at ~0 mW (4.2 s intervals, total 60 s). Dashed circles denote the laser focus of the trapping laser.

Supplementary Movie 13. Venus-SYP41 dynamics during trapping laser irradiation at 1 mW (4.2 s intervals, total 60 s). Dashed circles denote the laser focus of the trapping laser.

Supplementary Movie 14. Venus-SYP41 dynamics during trapping laser irradiation at 30 mW (4.2 s intervals, total 60 s). Dashed circles denote the laser focus of the trapping laser.

Supplementary Movie 15. Dynamics of vacuolar membranes and amyloplasts during trapping laser irradiation at 30 mW (2.1 s intervals, total 90 s). Dashed circles and white arrow denote the laser focus of the trapping laser and drastic deformation of the vacuolar membranes, respectively.

Supplementary Movie 16. Formation of transvacuolar strands by movements of trapped amyloplasts at 30 mW laser power and movements of amyloplasts after turning off the laser (2.1 s intervals, total 150 s). Dashed circles denote the laser focus of the trapping laser. White arrows in the left and right panel denote artificial formation of the transvacuolar strand and “spring back” response of the amyloplasts after switching off the trapping laser, respectively.

Supplementary Movie 17. Amyloplasts (yellow arrow) that moved away from the laser focus during trapping laser irradiation at 1 mW (2.1 s intervals, total 60 s). Dashed circles denote the laser focus of the trapping laser.

Supplementary Movie 18. Amyloplasts (yellow arrow) that moved away from the laser focus during trapping laser irradiation at 2 mW (2.1 s intervals, total 60 s). Dashed circles denote the laser focus of the trapping laser.

