## Supplementary figures and images for "Micromanipulation of amyloplasts with optical tweezers in *Arabidopsis* stems"

### Supplemental Figure 1

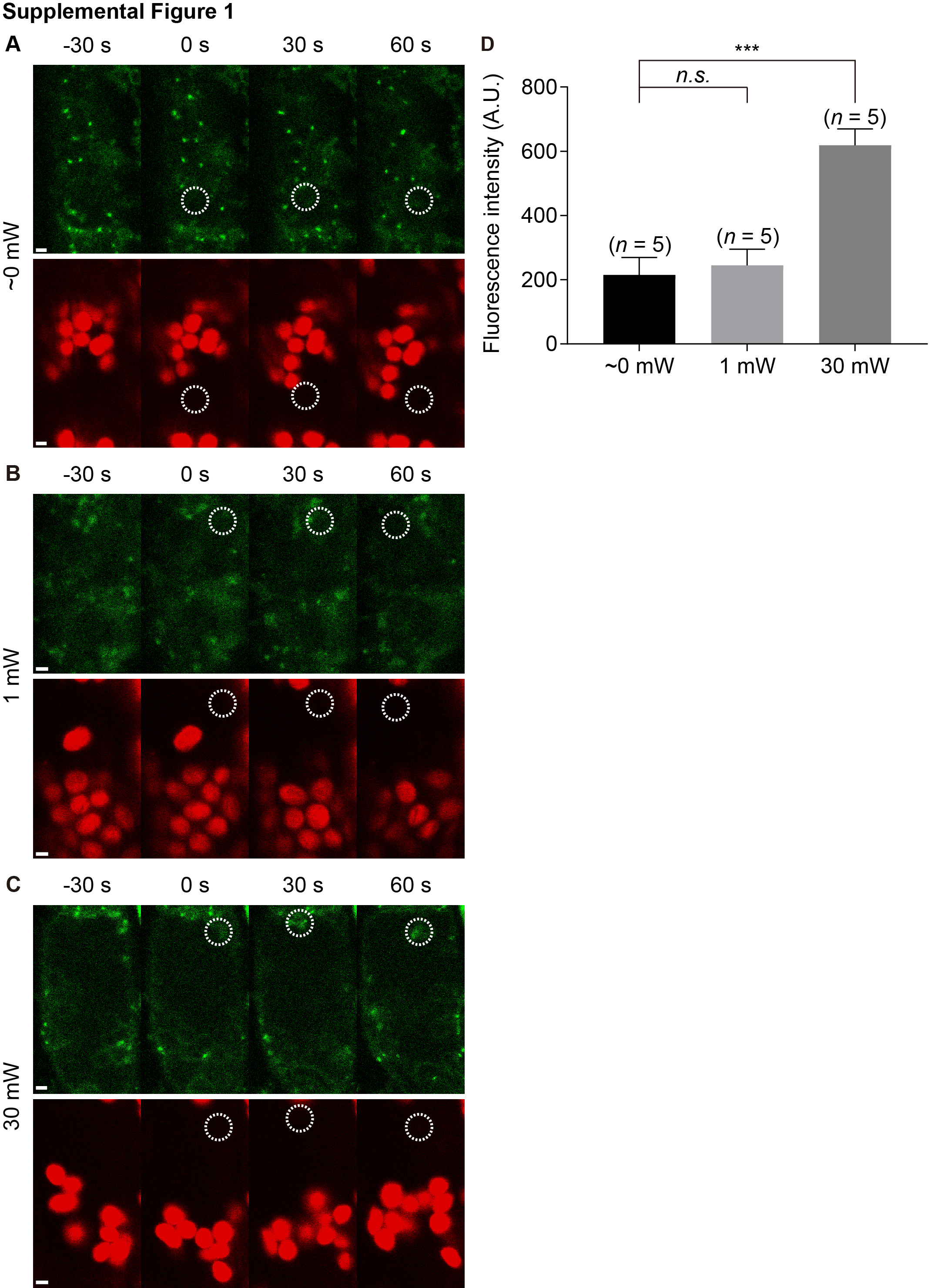

### Supplemental Figure 2

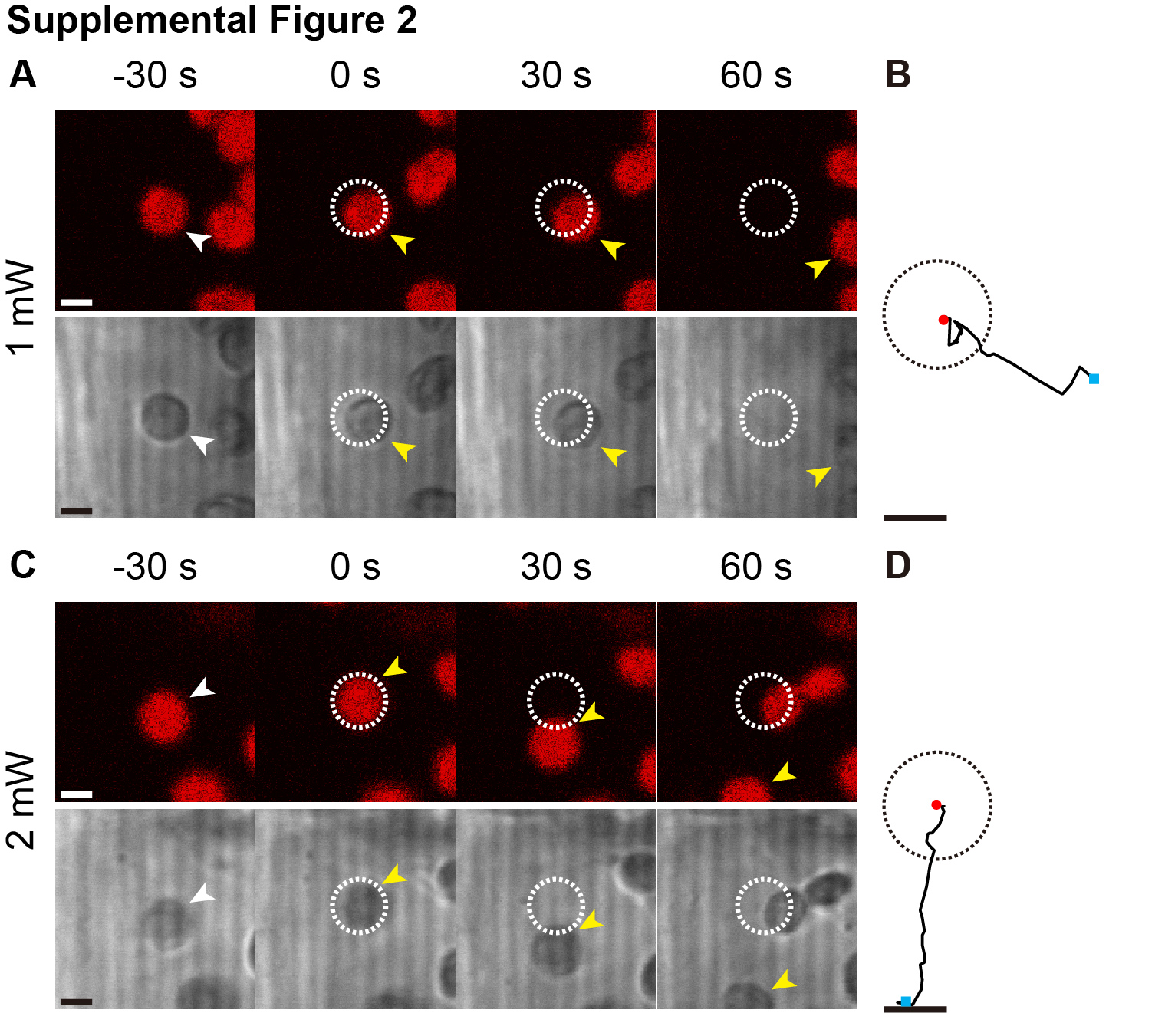
